# Influence of sequencing depth on bacterial classification and abundance in bacterial communities

**DOI:** 10.1101/2022.01.04.474922

**Authors:** Fernando Mejia Sanchez, Francisco Avilés Jiménez, Alfonso Méndez Tenorio

## Abstract

Microbial diversity is the most abundant form of life. Next Generation Sequencing technologies provide the capacity to study complex bacterial communities, in which the depth and the bioinformatic tools can influence the results. In this work we explored two different protocols for bacterial classification and abundance evaluation, using 10 bacterial genomes in a simulated sample at different sequencing. Protocol A consisted of metagenome assembly with Megahit and Ray Meta and taxonomic classification with Kraken2 and Centrifuge. In protocol B only taxonomic classification. In both protocols, rarefaction, relative abundance and beta diversity were analyzed. In the protocol A, Megahit had a mean contig length of 1,128 and Ray Meta de 8,893 nucleotides. The number of species correctly classified in all depth assays were 6 out of 10 for protocol A, and 9 out of 10 using protocol B. The rarefaction analysis showed an overestimation of the number of species in almost all assays regardless of the protocol, and the beta diversity analysis results indicated significant differences in all comparisons. Protocol A was more efficient for diversity analysis, while protocol B estimated a more precise relative abundance. Our results do not allow us to suggest an optimal sequencing depth at specie level.

## Introduction

Microbial diversity is composed of a great variety of unicellular organisms (prokaryota, archaea, protozoa, fungi and viruses) and is the most abundant life forms present on the planet [1]. Advances in next-generation sequencing (NGS) technologies have allowed us to reach unprecedented levels of genomic analysis [2], [3], and have provided us with the possibility to analyze non-culturable communities, whether they are animal tissue, air or soil samples [4].

Two sequencing methods are usually used in metagenomic studies, 16S and shotgun. The first focused in the sequencing of hypervariable regions of the 16S rRNA gene and the second consists of the sequencing of complete genomes from a sample [5]. Both methods allow the analysis of the alpha, beta and gamma diversity of a microbial community [6]. However, 16S sequencing has limitations, being a gene conserved in the prokaryote kingdom, its resolution is limited and it has a lower sensitivity in the identification of genus and species [7], [8]. The shotgun method increases resolution, sensitivity at the genus, species or bacterial strain level and bacterial co-abundance in microbiome studies [9]. However, generation of precise results depends not only on the sequencing platform, but also on the depth and data analysis employed [10]. It is important to highlight that there is not a consensus about the right protocol for processing and analysis of the sequencing data, due to the existence of different approaches and to the great variety of bioinformatics tools [8], [11].

In microbiome studies, the main bioinformatics toolsets are metagenome assemblers (MetasSPAdes, Megahit, Ray Meta) and taxonomic classifiers (Kraken and Centrifuge) that allow us to estimate the diversity and relative abundance of the microorganisms present in a sample [12]–[14].

The knowledge of the precise abundance at the genus and species level is important to better understand the ecological composition, the possible interactions, as well as the associations between pathology and specific microorganisms [15]. The aim of this work was to explore two different approaches for bacterial classification and abundance evaluation, using a simulated bacterial community sample at different sequencing depths.

## Materials and methods

### Computational resources

The present work was carried out on a server with Linux CentOS operating system, 35 processors and 62 GB RAM.

### Sample design

For the “in-silico” analysis, we used a simulated reference sample composed by 10 different bacterial genomes belonging to the main phyla that conform the human gut microbiota. Bacterial genomes were downloaded from the National Center for Biotechnology Information database (NCBI). To constitute the reference sample, we provided an arbitrary abundance for each one of the selected species, considering a total of 25 genomes which were distributed among the 10 species. In this context, the 25 genomes copies microbiome corresponds to the 1X depth for the reference sample. Table 1 shows the list of selected bacterial species, the number of copies of each genome, their genome size, abundance and the corresponded phylum.

**Table 1.**
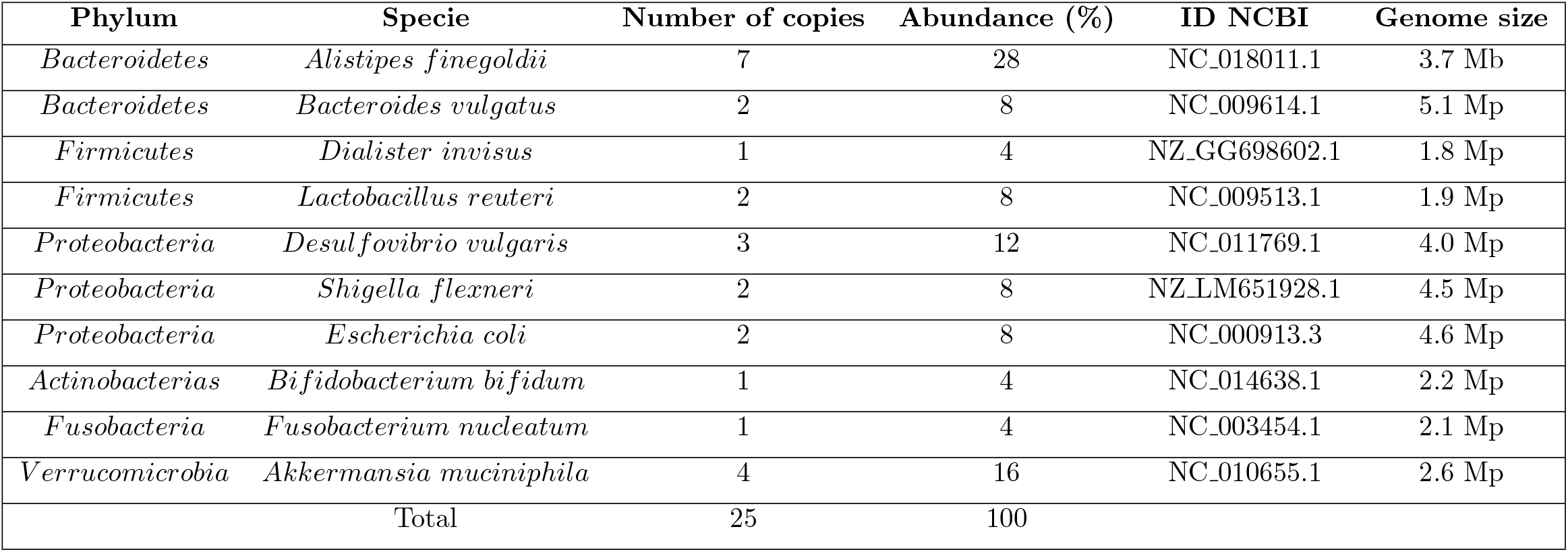
Species and bacterial abundance for the reference sample.

### Shotgun sequencing simulation

Using the reference sample, we simulated a shot-gun sequencing in silico for the Illumina HiSeq2500 platform and by means of the ART program [17] we obtained paired-end reads. In order to analyze the effect of sequencing depth on the bacterial community analysis, we generate 150mer reads for the following sequencing deeps of coverage: 1X, 25X, 50X, 75X, 100X, 125X and 150X.

### Sequence analysis

Taxonomic classification, rarefaction and bacterial abundance analyses were assessed by means of two sequence analysis protocols. Protocol A is divided into 4 stages: assembly, taxonomic classification and rarefaction, bacterial abundance, and statistical analysis, while protocol B includes all steps, but it does not consider the assembly step (Figure 1). Each de novo assembly was performed using a k-mer of 31, 51, 79 and 109 nucleotides.

**Fig 1.**
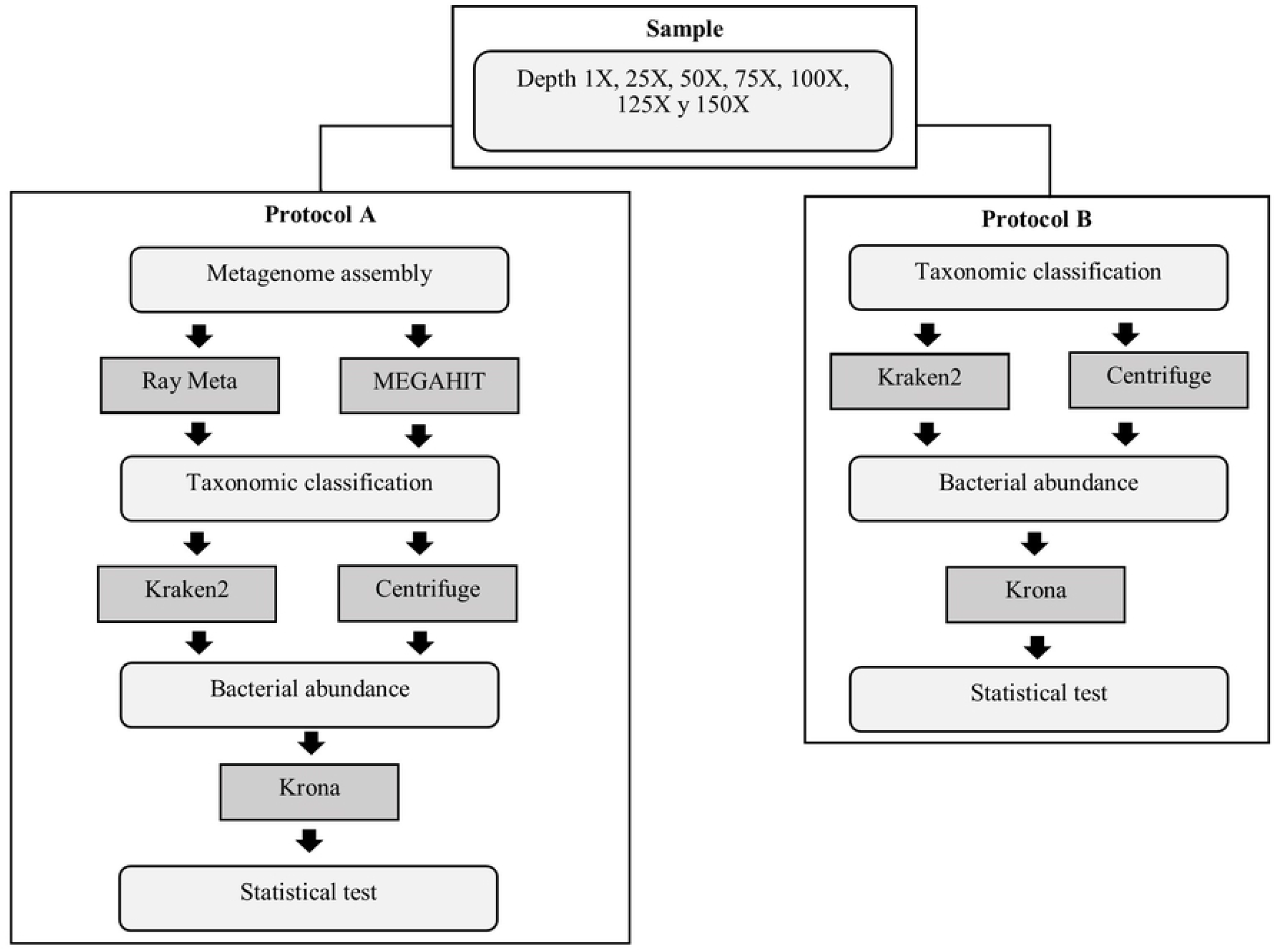
Protocols used in sequence analysis. Assembly was performed with MEGAHIT V 1.2.9 [4] and Ray Meta V 2.3.1 [16] programs, while bacterial classification and estimation of bacterial abundance was evaluated with Kraken2 V 2.1.2 [16] and Centrifuge V 1.0 [13]. Microbial-RefSeq and Bacteria-Archea databases were used respectively.

The classification results were obtained after filtering the information from each classifier considering only the information related to the 10 species that constitute the reference sample in each depth assay (Table 1). In contrast, for the analysis of total diversity, we included the whole number of species identified without any filtering step.

The estimated number of copies for each genome in the different depth assays was calculated by multiplying 25 (number of initial copies) by the depth value on each test. The relative abundance represents the number of copies of each species in relation to the total number of readings and expressed in percentage, for each one of the depth assays. Finally, the alpha diversity present in the reference and in each test was used for the calculation of beta diversity using the Sorensen index [18]. The statistical comparison of species between the reference sample and each depth assay was carried out by means analysis of an analysis of variance (ANOVA).

## Results

### Metagenome assembly

In the assembly stage (protocol A), Ray Meta had better results than Megahit if we consider the number and size of the assemblies. Ray Meta generated a lower number of contigs, and at the same time those had a larger size. The average length was 8,893 nucleotides with Ray Meta and 1,128 nucleotides with Megahit (Table 2).

**Table 2.**
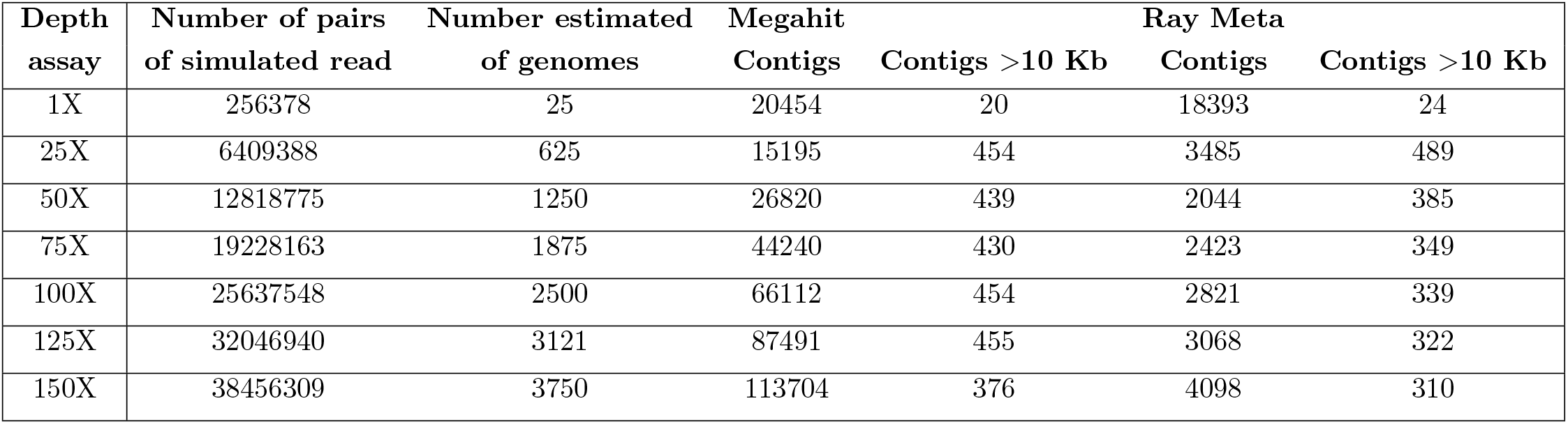
Number of reads, genomes and contigs.

| Depth assay | Number of pairs of simulated read | Number estimated of genomes | Megahit |  | Ray Meta |  |
| --- | --- | --- | --- | --- | --- | --- |
|  |  |  | Contigs | Contigs >10 Kb | Contigs | Contigs >10 Kb |
| 1X | 256378 | 25 | 20454 | 20 | 18393 | 24 |
| 25X | 6409388 | 625 | 15195 | 454 | 3485 | 489 |
| 50X | 12818775 | 1250 | 26820 | 439 | 2044 | 385 |
| 75X | 19228163 | 1875 | 44240 | 430 | 2423 | 349 |
| 100X | 25637548 | 2500 | 66112 | 454 | 2821 | 339 |
| 125X | 32046940 | 3121 | 87491 | 455 | 3068 | 322 |
| 150X | 38456309 | 3750 | 113704 | 376 | 4098 | 310 |

### Taxonomic classification

The taxonomic classification in protocol A was equally efficient when Centrifuge or Kraken2 were used. In the protocol A, we evaluated two different assemblers, the first assembler was Megahit combined with two different classifiers. At this stage, both the Centrifuge and Kraken2 classified 6 of the 10 species that conform the reference sample, and the remaining four species were classified intermittently or were absent. The same results were observed repeatedly in all depth assays (Figure 2a and Figure 2b).

**Fig 2.**
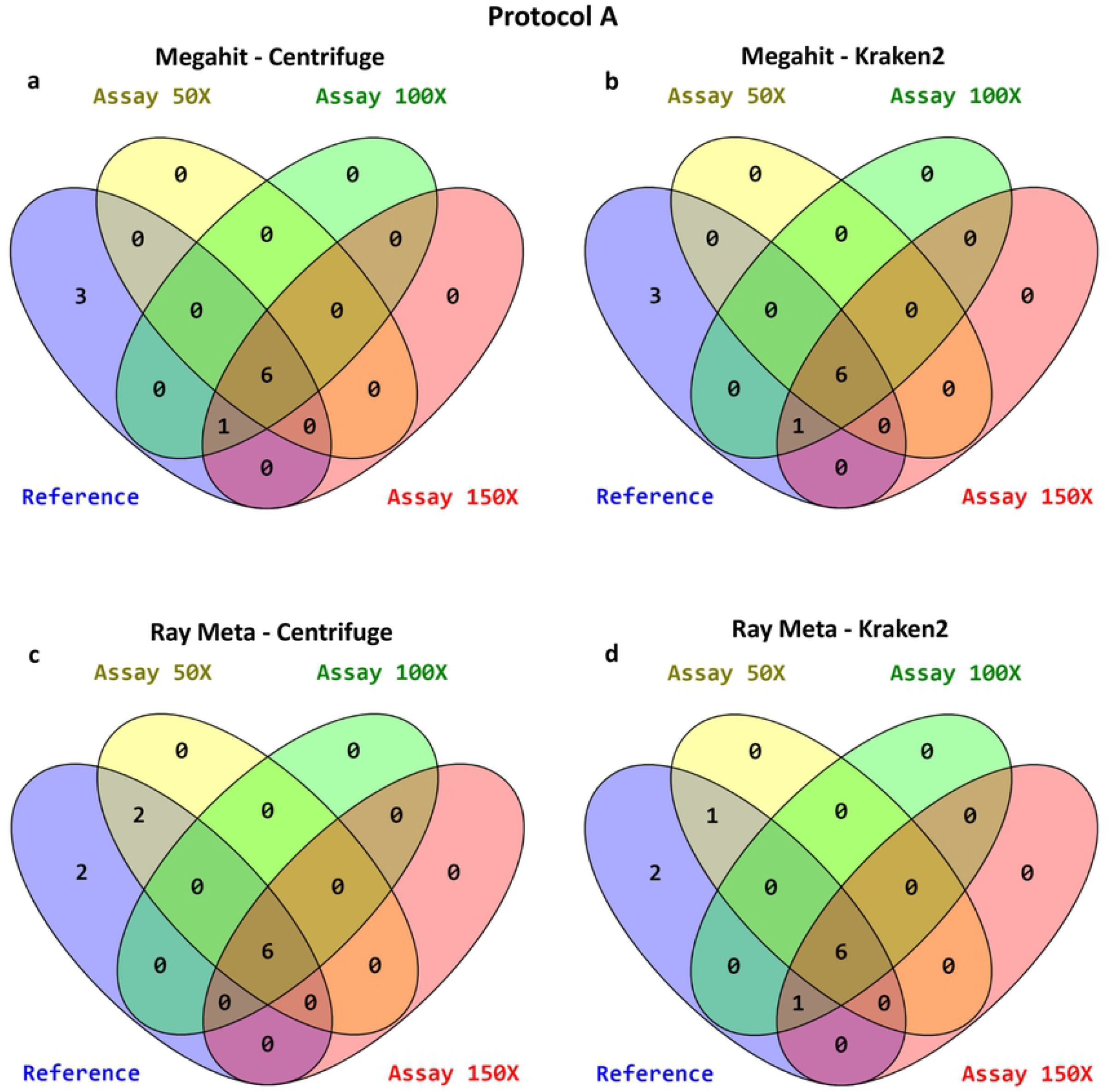
Species of interest classified using the protocol A. Venn diagrams showing the number of species classified for each one of the depth assays, according to the following color code: Blue (Reference), Yellow (50X), Green (100X) and Red (150X). In all assays, 6 out of the 10 species from the reference sample were consistently well classified in all assays (*Akkermansia muciniphila*, *Alistipes finegoldii*, *Bacteroides vulgatus*, *Escherichia coli*, *Lactobacillus reuteri* and *Shigella flexneri*). The remaining four were classified intermittently or were not identified (*Bifidobacterium bifidum*, *Desulfovibrio vulgaris*, *Dialister invisus* and *Fusobacterium nucleatum*). a: Megahit assembler combined with Centrifuge classifier. b: Megahit assembler combined with Kraken2 classifier. c: Ray Meta assembler combined with Centrifuge classifier. d: Ray Meta assembler combined with Kraken2 classifier.

Using Ray Meta assembler, we obtained similar findings, 6 out of 10 species were properly classified with either Centrifuge and Kraken 2 classifiers in all depth assays (Figure 2c and 2d).

Taxonomic classification without previous assembly (protocol B) yielded more efficient results. In this protocol we only evaluated the classifiers Centrifuge and Kraken2, both classified 9 of the 10 species present in the reference sample and, the remaining specie was not classified in any assay. These results were consistent in all depth assays (Figure 3a and 3b) and, shows the same level of efficiency of both taxonomic classifiers.

**Fig 3.**
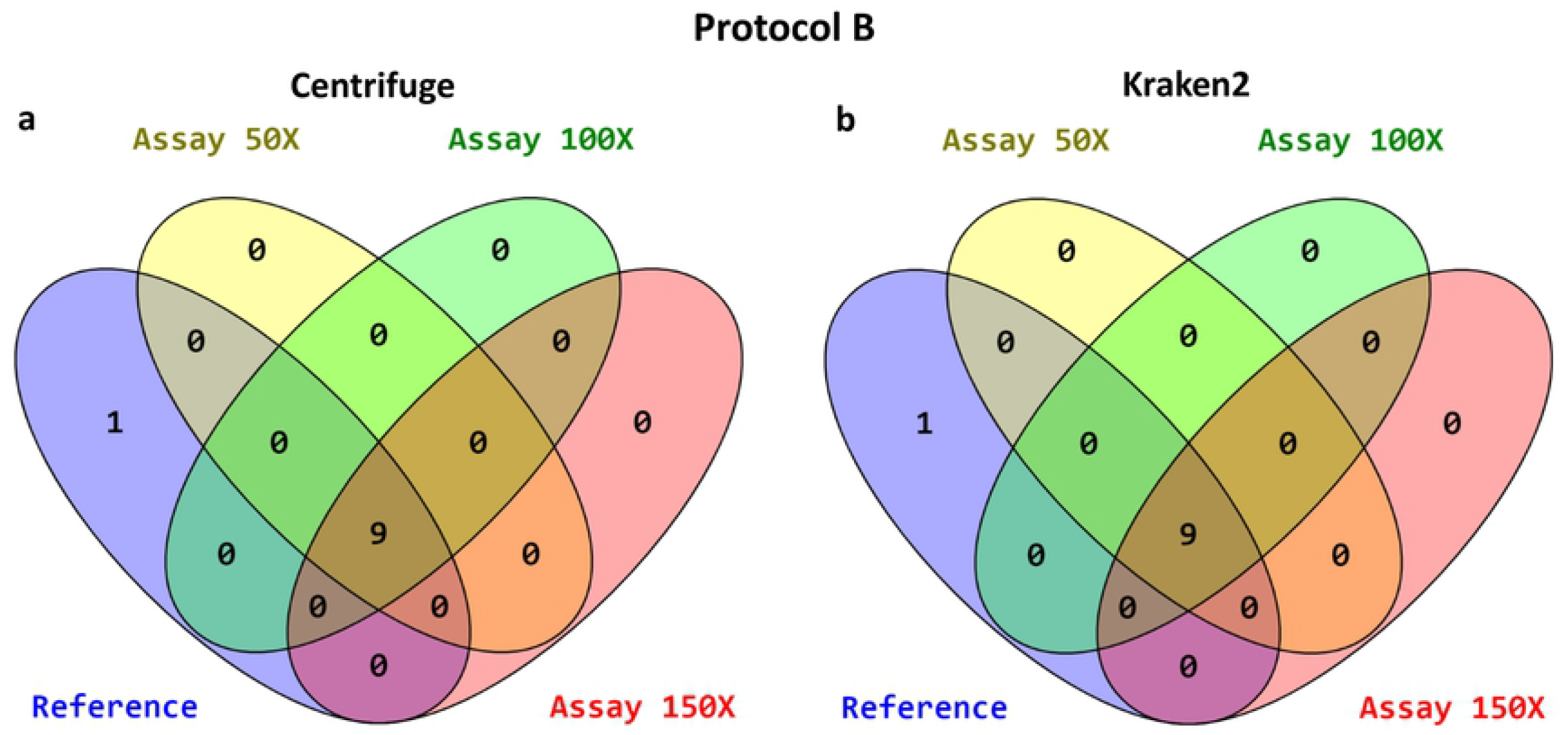
Species of interest classified using the protocol B. Venn diagrams showing the number of species classified for each one of the depth assays, according to the following color code: Blue (Reference), Yellow (50X), Green (100X) and Red (150X). In all assays, 9 out of the 10 species from the reference sample were consistently well classified in all assays (*Akkermansia muciniphila*, *Alistipes finegoldii*, *Bacteroides vulgatus*, *Bifidobacterium bifidum*, *Desulfovibrio vulgaris*, *Escherichia coli*, *Fusobacterium nucleatum*, *Lactobacillus reuteri* and *Shigella flexneri*). The remaining specie was not classified (*Dialister invisus*). a: Centrifuge classifier. b: Kraken2 classifier.

### Rarefaction analysis

The determination of diversity and species richness were more accurate when a prior assembly process is implemented. In the protocol A, the Megahit assembler combined with each of the two classifiers (Centrifugue and Kraken2) recorded a higher number of species compared to the Ray Meta assembler, and this result was ascending in each depth assay. In both cases, the richness was greater than that present in the reference sample (Table 3 and Figure 4).

**Table 3.** Total number of species of the protocol A and B.

| Depth assay | Protocol A |  |  |  | Protocol B |  |
| --- | --- | --- | --- | --- | --- | --- |
|  | Megahit |  | Ray Meta |  | Centrifuge | Kraken2 |
|  | Centrifuge | Kraken2 | Centrifuge | Kraken2 |  |  |
| 1X | 51 | 68 | 26 | 12 | 446 | 304 |
| 25X | 105 | 91 | 192 | 108 | 1919 | 1225 |
| 50X | 140 | 112 | 29 | 16 | 2181 | 1663 |
| 75X | 190 | 159 | 21 | 14 | 2285 | 1964 |
| 100X | 222 | 186 | 24 | 16 | 2344 | 2240 |
| 125X | 218 | 194 | 21 | 15 | 2395 | 2466 |
| 150X | 251 | 221 | 22 | 17 | 2457 | 2559 |

**Fig 4.**
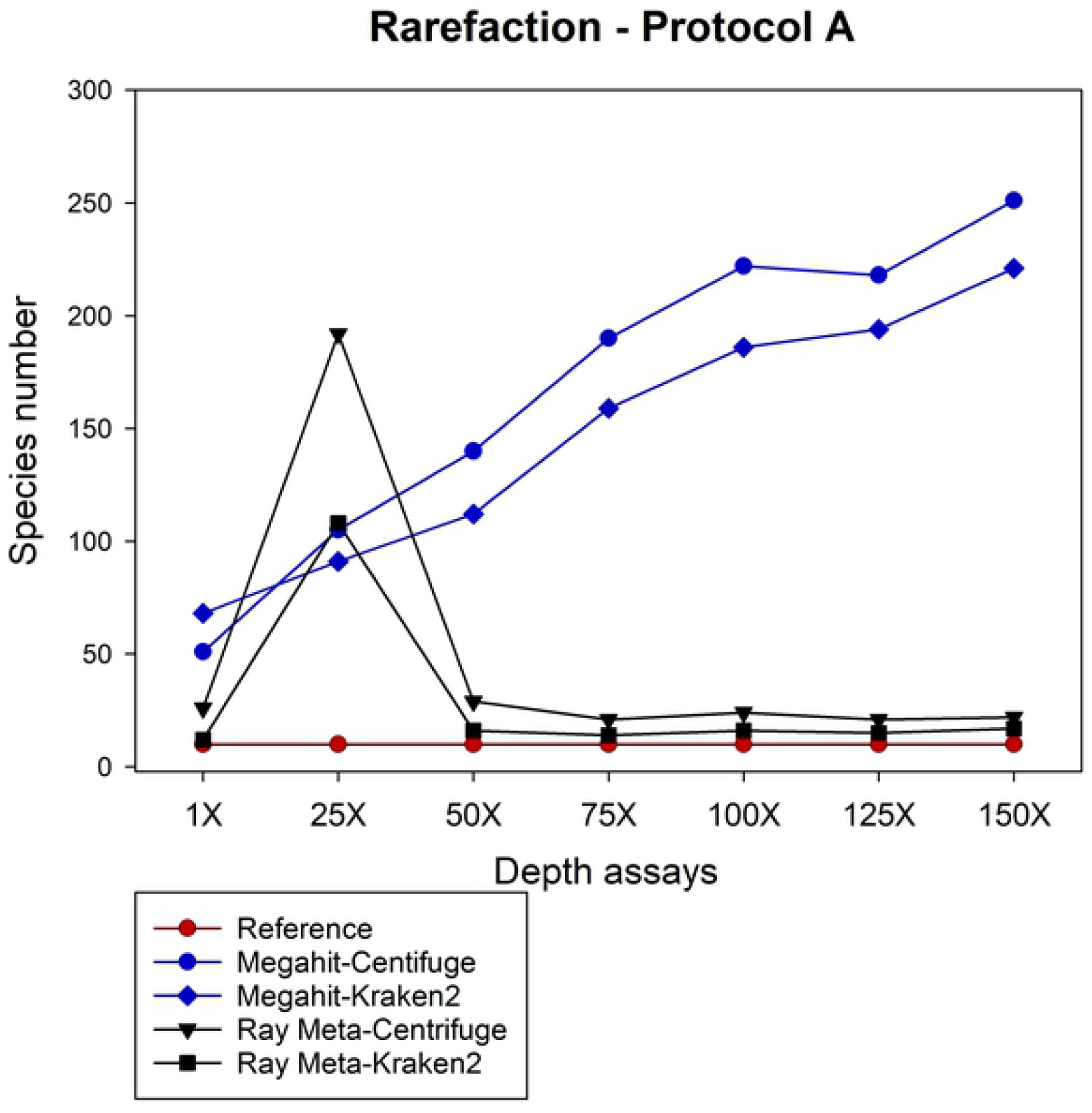
Rarefaction curves showing the total number of identified species using the protocol A. The number of species identified in each depth assay was greater than the theoretical value of the reference sample. Of the two assemblers used, Megahit registers a greater diversity with respect to that reported by Ray Meta for both classifiers. For the specific number of species identified see Table 3.

On the other hand, when the data are analyzed with protocol B without including a previous assembly process, the total number of species identified, considerably exceeds the theoretical value of the reference sample in each depth assay, regardless of the classifier used.(Table 3 and Figure 5).

**Fig 5.**
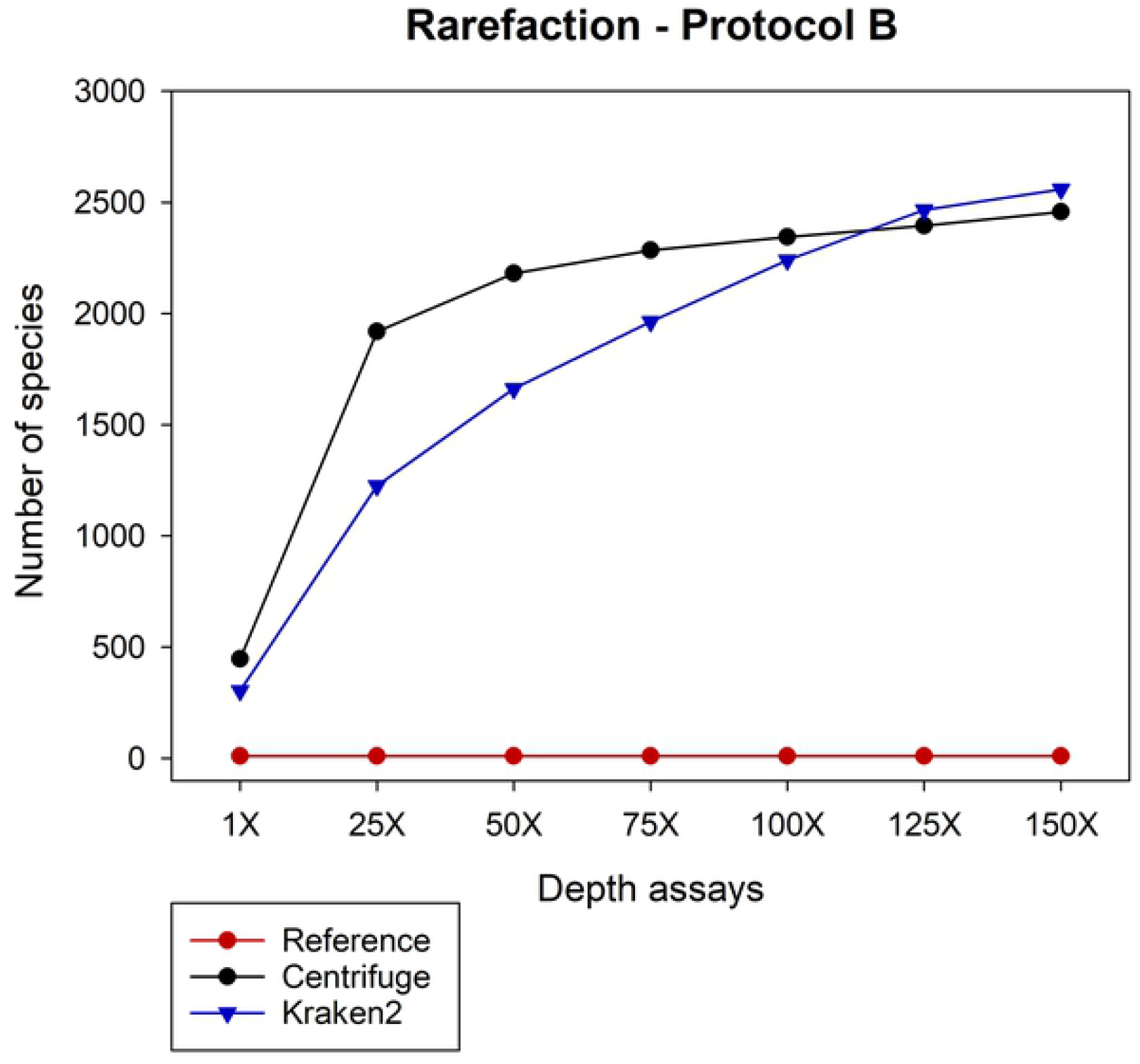
Rarefaction curves for total number of identified species using the protocol B. The number of species identified with protocol B is much higher than the number of species reported for protocol A. This growth pattern is maintained for each one of the depth assays and holds the same trend with any of the two classifiers. For the specific number of species identified see Table 3.

### Relative abundance

The estimation of the relative abundance of the bacterial species classified in the assemblies was inaccurate and this is visible with Megahit and Ray Meta. Addressing protocol A, the relative abundances estimated with Centrifuge and Kraken2 in the Megahit assemblies differed than that present in the reference sample, these results were observed in all depth assays (Figure 6). On the other hand, classification of Ray Meta assemblies generated the most imprecise relative abundances regardless of taxonomic classifier (Figure 7).

**Fig 6.**
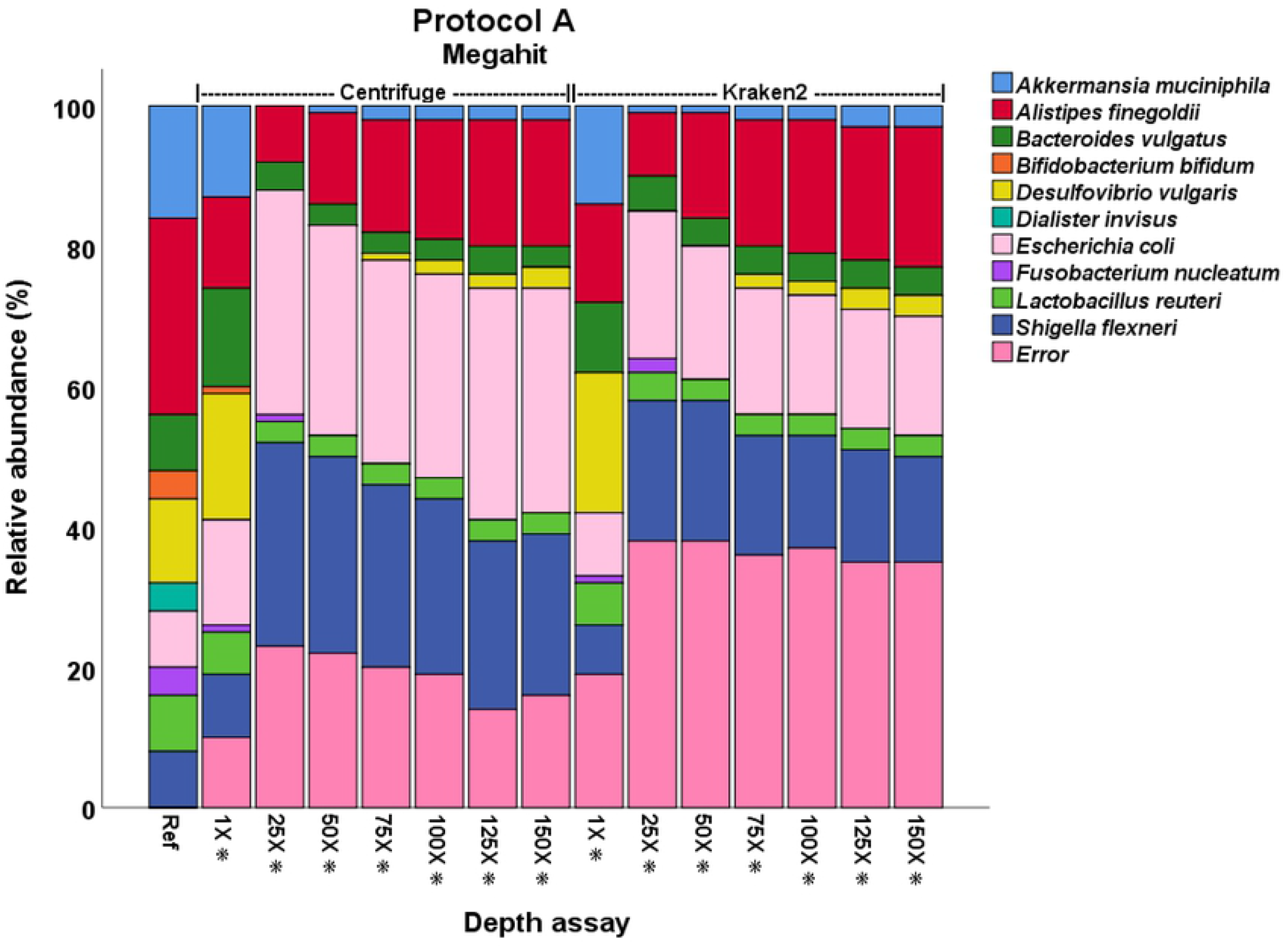
Relative abundance in Megahit assemblies of protocol A. Kraken2 generated a higher abundance of classification errors. *Significant difference in the analysis of beta diversity between depth assays and the reference, p<0.005.

**Fig 7.**
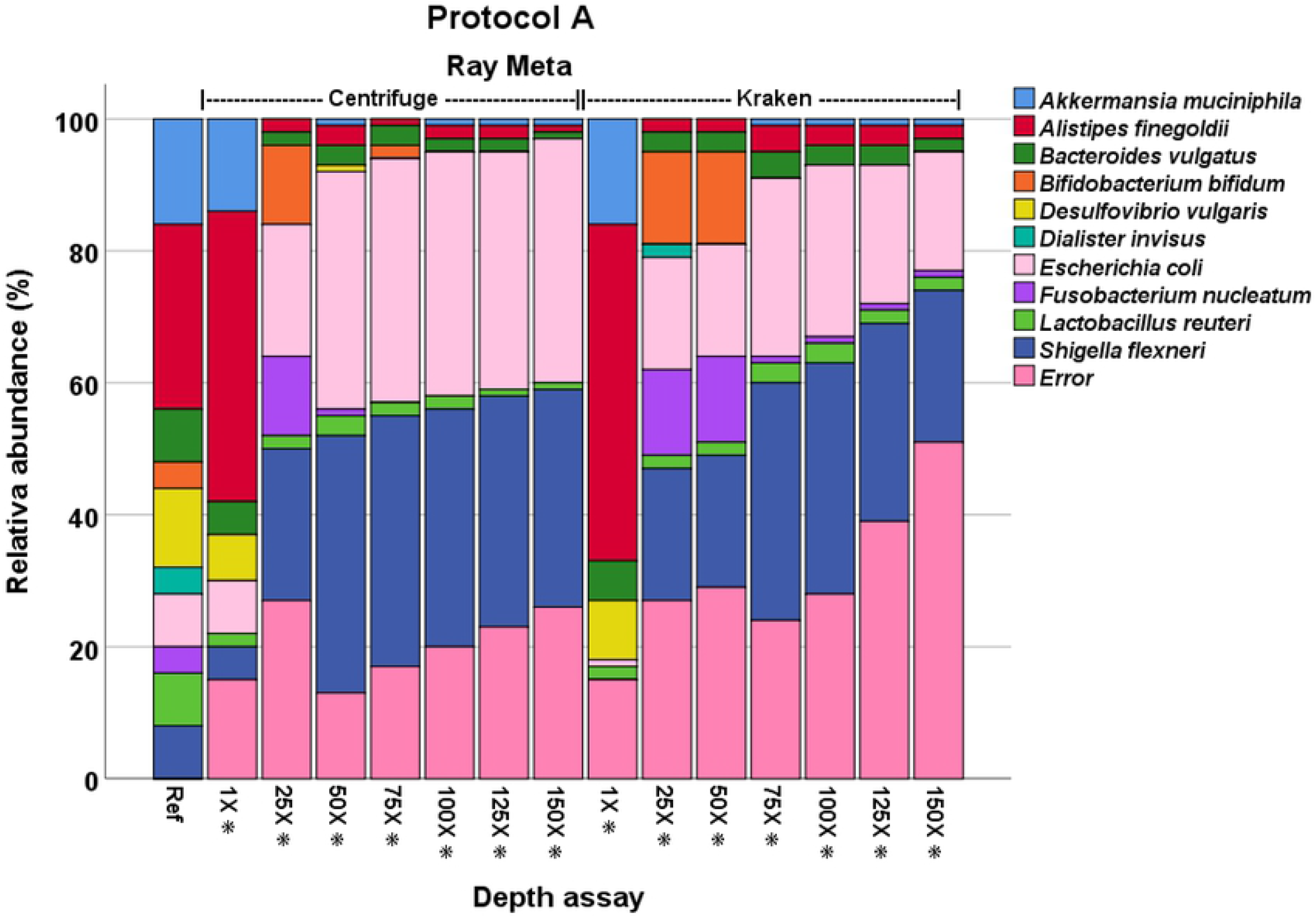
Relative abundance of species in Ray Meta assemblies of protocol A. Kraken2 generated a higher abundance of classification errors. +Significant difference in the analysis of beta diversity between depth assays and the reference, p<0.005.

By excluding assemblages and using protocol B, the estimation of the relative abundance of bacterial species produced better results. In this protocol, Centrifuge and Kraken2 generated similar relative abundances in relation to the reference and this result was obtained repeatedly in all assays (Figure 8).

**Fig 8.**
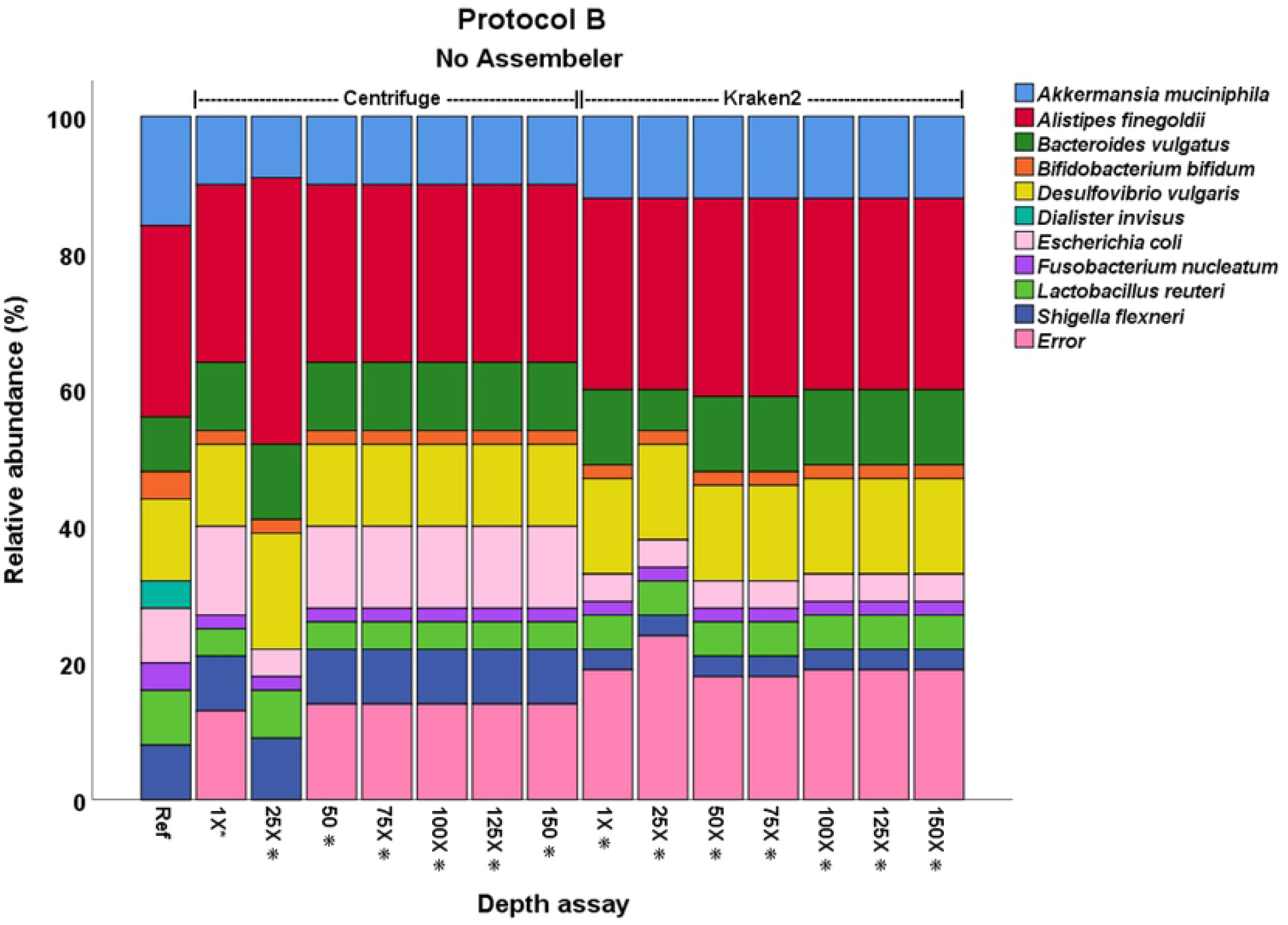
Relative abundance of species using protocol B. Relative abundances exhibit minor variations between the different depth assays with any of the two classifiers. *Significant difference in the analysis of beta diversity between depth assays and the reference, p<0.005.

### Beta diversity analysis

The beta diversity analysis at the species level between the different depth assays and the reference in protocol A, showed significant differences (p<0.005). (Figures 6 and 7). The same result was obtained for the protocol B (p<0.005). (Figure 8). When analyzing the beta diversity between each one of the depth assays with all the others we observed a significant difference (p<0.005).

In the protocol A, at the genus level, the use of the centrifuge classifier with Megahit or Ray Meta assemblers, showed no significant difference from the 50X depth assay and the results did not improve after this point (p>0.005). In contrast, using protocol B, there was no significant differences from the 25X assay with Centrifuge (p>0.005).

## Discussion

This work has focused on exploring some of the tools commonly used in the characterization of microbial communities using shotgun sequencing and that do not require extensive knowledge or abilities in bioinformatics [19], [20]. Considering this, in this work we did not explore in depth the mathematical basis or logarithms of these tools. The use of a simulated sample in this work helps to eliminate the negative influence of errors or low sequencing quality of a real sample [21]. In this study, we determined the diversity and abundance in each assay, this gives greater certainty in the results obtained.

The results of the metagenome assemblies showed that about 90 of the contigs had a size below 10,000 nucleotides in the protocol A. This size is smaller than the genome of *Dialister invisus* (1.8 Mb), specie in the sample with the smaller genome size. In this work we used Megahit and Ray Meta, these are “de novo” assemblers that are both based on Bruijn graphs [4], [16]. This type of graphs allows the efficient assembly of short reads [22], but when divided into the length of the k-mer defined in the assembly, these can be susceptible to errors [12]. During the assembly process of a mixed genome, two types of errors often occur. The first is the presence of k-mer in different regions of the same genome, giving rise to chimeric connections between the nodes in the Bruijn graph, which are different to the real sequence. This can result in erroneous assemblies and short contigs. This error increases when assembly a metagenome because a k-mer can be present in different genomes. The second error occurs in regions with low coverage. However, this error was excluded when a simulated sample was studied [23]. The presence of short contigs in this study is the result of erroneous assemblies and the use of k-mer of different lengths did not prevent the generation of these errors. Currently, the use of next-generation sequencing platforms such as PacBio and Nanopore, which can sequence fragments of 30-50 Kb and 100 Kb respectively [24]. This may be favorable in the study of complex bacterial communities, because larger read sizes can generate more precise assemblies, even with shared genomic regions.

Regarding the efficiency of the assemblers, there are very few publications where a comparison of these tools is performed. The reports that introduced Ray Meta and Megahit, based the results of assembler efficiency on the comparison of their results with respect to other assemblers, and they gave greater relevance to avoidance of the assembly of repeated reads, ramifications and the generation of longer assemblies. However, these parameters are not a guaranty of their efficiency, which we observed in the assemblies performed in this study [4], [12], [16]. In addition, the performance evaluation was not carried with a sample that has a known composition.

The classification of the assemblages showed a lower number of species than the number of contigs obtained in each assay, this was due to the absence of genomic information necessary for their classification. In contrast, the number of species was overestimated in each of the assays, and this influenced the beta diversity and relative abundance, which were different from the reference. The results obtained in this work related to the assemblages, show the need to discriminate the short assemblages, since there may be misclassified and influence the results of rarefaction, relative abundance, and beta diversity.

The omission of metagenome assembly and the exclusive use of taxonomic classifiers is common in microbiota studies where shotgun sequencing is used [10], [25], this can be explained by the high demand of computational requirements and processing time of the assemblers [25]. Our results from protocol B reflect a relative abundance different to the reference and overestimation of species much higher than protocol A in each of the depth assays. This is of utmost importance since bacterial richness and relative abundance are two of the most important results in microbiota studies.

The results obtained with both protocols indicate that neither of the bioinformatics tools used in this study is completely accurate, but Ray Meta generated the larger size assemblies and avoided the generation of short contigs. While Centrifuge had the lowest number of errors, but this was influenced by the type of database, which includes genomic information characteristic of bacterial species and omits redundancies [13]. This makes the classification of species with shared genomic regions more efficient.

Regarding the influence of depth, we are not able to obtain information on the optimal metagenome sequencing depth to obtain reliable taxonomic results. However, at the genus level, we did not observe an improvement in the results from the 50X depth and using Ray Meta with Centrifuge.

It is important to note that the sample analyzed in this study is minimal and cannot be compared to the composition of a real complex bacterial community sample. Nevertheless, the information generated can help researchers in the selection of bioinformatics tools and the use of data filtering strategies for metagenome assemblies and taxonomic classification.

## Conclusion

For metagenome assembly and diversity analysis, protocol A (Ray Meta and Centrifuge) was more efficient. While in relative abundance, protocol B (Centrifuge) was better. Our results do not allow us to suggest an optimal sequencing depth.

